# Recapitulation of SARS-CoV-2 Infection and Cholangiocyte Damage with Human Liver Organoids

**DOI:** 10.1101/2020.03.16.990317

**Authors:** Bing Zhao, Chao Ni, Ran Gao, Yuyan Wang, Li Yang, Jinsong Wei, Ting Lv, Jianqing Liang, Qisheng Zhang, Wei Xu, Youhua Xie, Xiaoyue Wang, Zhenghong Yuan, Junbo Liang, Rong Zhang, Xinhua Lin

## Abstract

The newly emerged pandemic coronavirus, SARS-CoV-2, has posed a significant public health threat worldwide. However, the mode of virus transmission and tissue tropism is not well established yet. Recent findings of substantial liver damage in patients and ACE2+ cholangiocytes in healthy liver tissues prompted us to hypothesize that human liver ductal organoids could serve as a model to determine the susceptibility and mechanisms underlining the liver damage upon SARS-CoV-2 infection. By single-cell RNA sequencing, we found that long-term liver ductal organoid culture preserved the human specific ACE2+ population of cholangiocytes. Moreover, human liver ductal organoids were permissive to SARS-CoV-2 infection and support robust replication. Notably, virus infection impaired the barrier and bile acid transporting functions of cholangiocytes through dysregulation of genes involved in tight junction formation and bile acid transportation, which could explain the bile acid accumulation and consequent liver damage in patients. These results indicate that control of liver damage caused directly by viral infection should be valued in treating COVID-19 patients. Our findings also provide an application of human organoids in investigating the tropism and pathogenesis of SARS-CoV-2, which would facilitate novel drug discovery.

## Introduction

A recent outbreak of SARS-CoV-2 (previously named 2019-nCoV) infection in Wuhan (China) has caused emergent and significant threats to global public health^1^. The dominant symptoms of coronavirus disease 2019 (COVID-19) are fever and cough^2,3^. However, a proportion of patients showed multi-organ damage and dysfunction^2-4^. Of note, liver damage is emerging as a co-existed symptom reported in patients with COVID-19. A recent epidemiologic study in Shanghai (China) reported that 75 out of 148 (50.7%) COVID-19 patients had liver function abnormality, indicated by key liver function parameters above the normal range, including alanine aminotransferase (ALT), aspartate aminotransferase (AST), alkaline phosphatase (ALP) or total bilirubin (TBIL)^5^. A national wide clinical study collecting 1,099 COVID-19 patients revealed that around 20% of patients had elevated ALT and AST and around 10% of patients had elevated TBIL. Especially, the percentage of patients with liver damage is much higher in severe patients than that in non-severe ones^2^. Although the clinical correlation has been implicated, it is still unclear whether the liver damage is caused by direct virus infection in the liver or by systematic reasons such as cytokine storm.

Viruses bind to host receptors to initiate the infection. Recent studies have demonstrated that both SARS-CoV-2 and SARS-CoV use the same angiotensin-converting enzyme 2 (ACE2) protein to enter the cells^6-10^. It has been shown that ACE2 expression is widely distributed across human tissues, including lung, liver, kidney and multiple digestive tract organs^11-13^. A significant enrichment of ACE2+ population in cholangiocytes compared to hepatocytes in human healthy liver was reported recently^14^, implying that SARS-CoV-2 might directly target ACE2+ cholangiocytes in patients. However, whether the virus indeed infects human cholangiocytes thus causes local damage has not been addressed yet.

At present, due to the lack of suitable research models, studies on mechanisms of SARS-CoV-2 pathogenesis mainly depend on bioinformatics analysis, clinical characteristics and rare autopsy reports^15^. Here we report the use of human organoids as a tool to investigate the SARS-CoV-2 infection and induced tissue damage ex vivo at the cellular and molecular levels. By single-cell RNA sequencing, we found that long-term human liver ductal organoid culture preserved the human specific ACE2+ population of cholangiocytes. Moreover, human liver ductal organoids were susceptible to SARS-CoV-2 infection and support robust viral replication. Notably, virus infection impaired the barrier and bile acid transporting functions of cholangiocytes in human liver ductal organoids. These results suggest that the dysfunction of cholangiocytes induced by SARS-CoV-2 infection could contribute to the bile acid accumulation and consequent liver damage in patients, and control of liver damage should be valued in treating COVID-19 patients. Our findings also provide a useful model of SARS-CoV-2 infection for pathogenesis study and novel drug discovery.

## Results

### ACE2+ cholangiocytes are preserved in human liver ductal organoid cultures

To establish the SARS-CoV-2 infection model with human liver ductal organoids, we first determined whether the long-term organoid culture could preserve the ACE2+ cholangiocytes ex vivo. We processed single-cell RNA sequencing (scRNA-seq) to interrogate the transcriptome of cholangiocytes in human liver ductal organoids. A total number of 7,978 cells were analyzed and cell populations were visualized by t-distributed stochastic neighbor embedding (t-SNE), partitioning the cells into 7 clusters (Fig. 1A). The common cholangiocyte markers *EPCAM* and *KRT19* were uniformly highly expressed in all the 7 clusters, indicating the heterogeneity of cholangiocytes in these organoids was relatively low (Fig. 1B). Notably, we identified the SARS-CoV-2 receptor gene *ACE2* expressed sparsely among cluster 0-5 in unbiased preferences and was detectable in 2.92% cells (233 out of 7,978) (Fig. 1C, D). Anti-ACE2 immunostaining further verified the presence of ACE2+ cholangiocytes in human liver ductal organoids (Fig. 1E). Interestingly, we found that the cholangiocytes in mouse primary liver ductal organoids had comparable *Epcam* expression but no *Ace2* (mouse *Ace2*) expression (Fig. 1F). Taken together, our data demonstrate that long-term human liver ductal organoid culture preserves the human specific ACE2+ population of cholangiocytes and the human liver ductal organoids could serve as a model to study the SARS-CoV-2 infection mediated by receptor ACE2.

**Figure 1.**
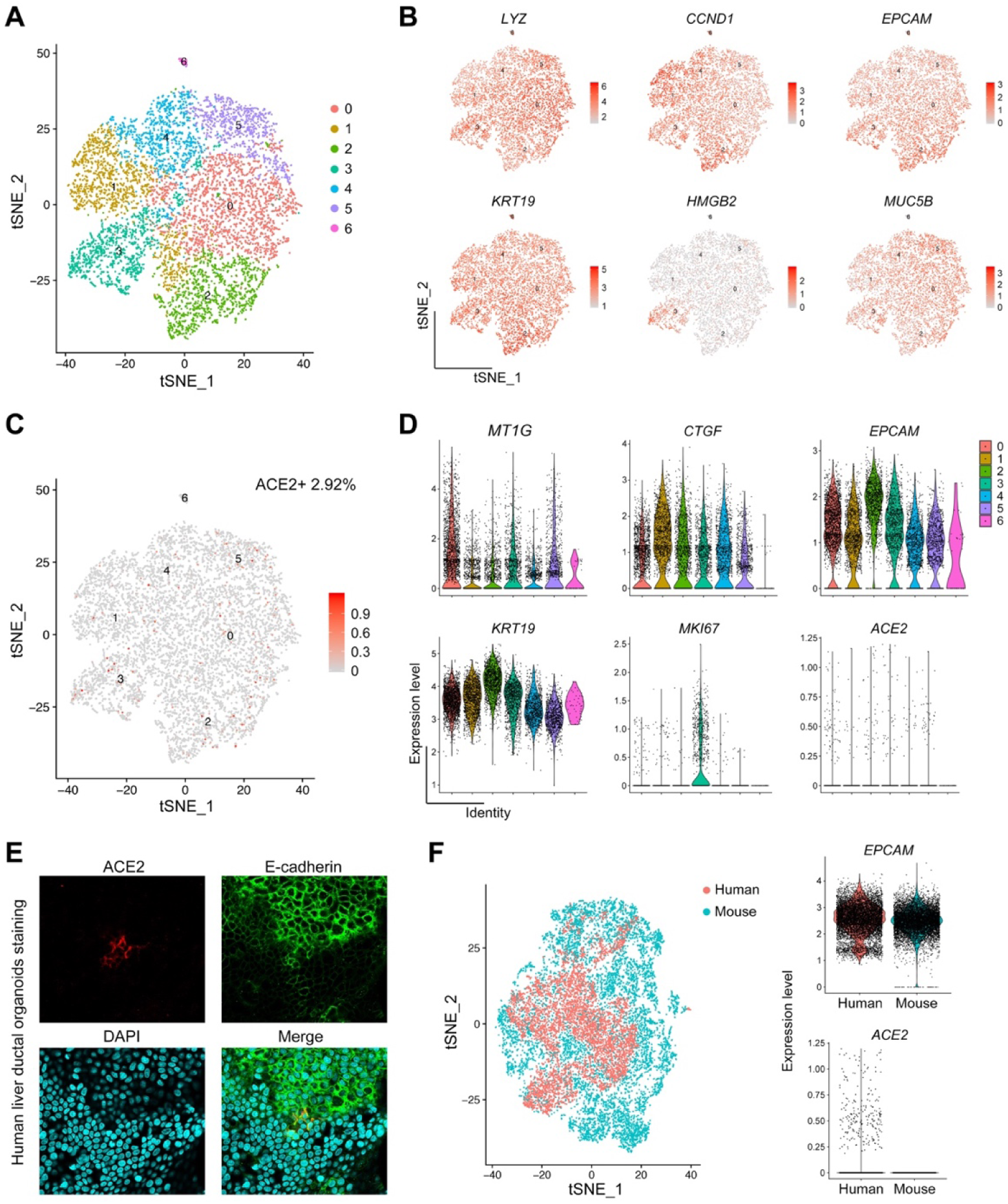
ACE2+ cholangiocytes are preserved in human liver ductal organoid cultures. (**A**) Cell-type clusters. t-SNE visualization of the cell populations (color-coded for clusters) from human liver ductal organoids by t-SNE. (**B**) t-SNE plots indicating the expression of representative marker genes. (**C**) t-SNE plots indicating the expression of ACE2 gene. (**D**) Violin plots showing the expression of representative marker genes. (**E**) Immunofluorescence staining for ACE2 and E-cadherin in human liver ductal organoids. Results were representative of three independent experiments. (**F**) t-SNE visualization of single cells isolated from human and mouse liver ductal organoids; Violin plots showing the expression of EPCAM and ACE2.

### Recapitulation of SARS-CoV-2 infection in human Liver ductal organoids

We next examined the susceptibility of human liver ductal organoids to SARS-CoV-2. We isolated and plaque-purified the SARS-CoV-2 from a COVID-19 patient in Shanghai. The liver ductal organoids from two individuals were inoculated with SARS-CoV-2 for 1 hour then re-embedded in Matrigel and maintained in culture medium. We performed immunostaining to identify the virus-positive cholangiocytes 24 hours post infection. The expression of SARS-CoV-2 nucleocapsid protein was readily detected in patchy areas of human liver ductal organoids whereas no signal was found in uninfected control (Figure 2A). In addition, the infected cholangiocytes underwent membrane fusion and formed syncytia (Figure 2A, enlarge). Although the number of infected cholangiocytes was limited, qRT-PCR analysis of the SARS-CoV-2 genomic RNAs revealed a dramatic increase of viral load in organoids at 24 hours post infection (Figure 2B). These data demonstrate that human liver ductal organoids are susceptible to SARS-CoV-2 and support robust viral replication. The recapitulation of SARS-CoV-2 infection in human organoids suggests that this model could be employed to dissect the viral pathogenesis and to test antivirals.

**Figure 2.**
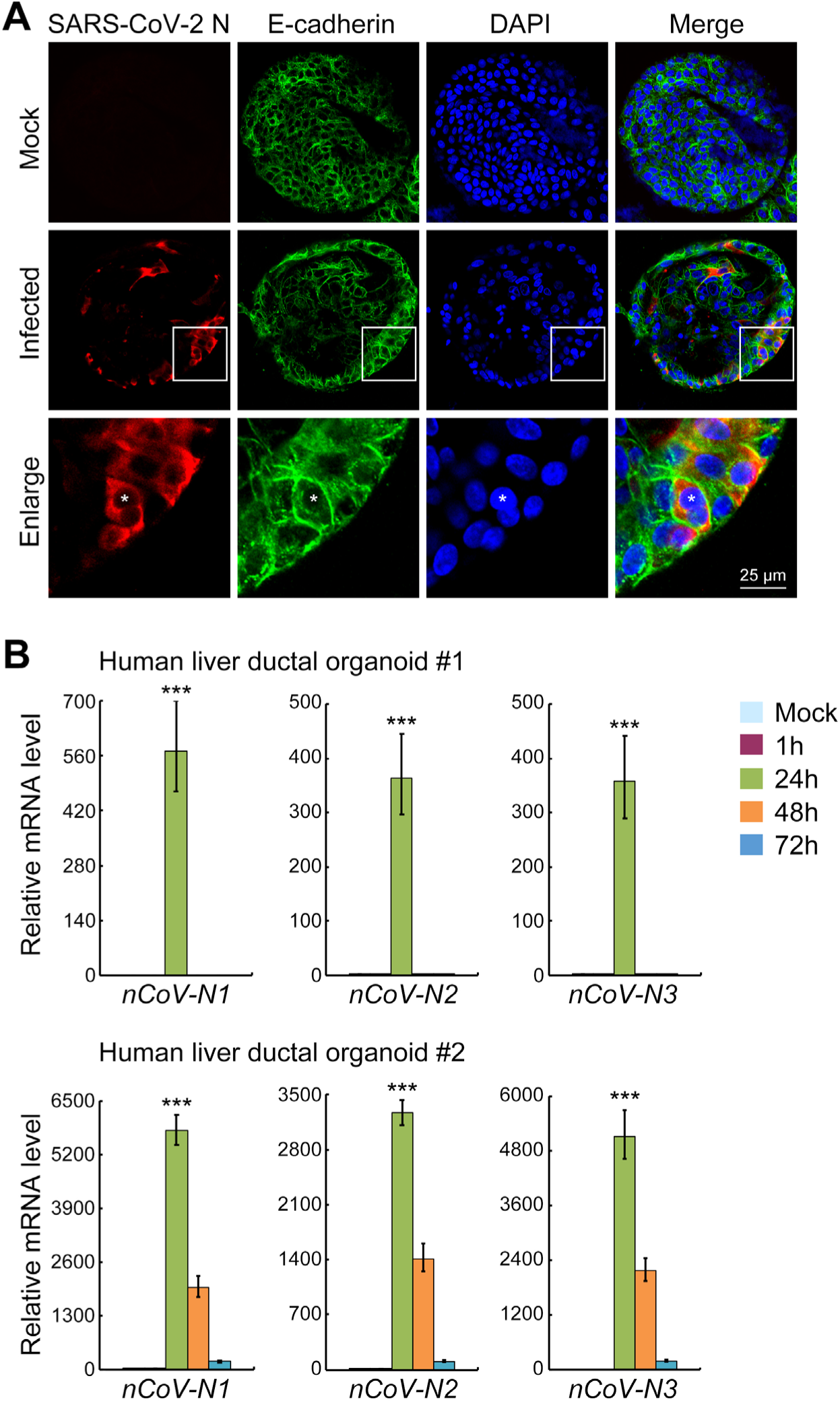
Recapitulation of SARS-CoV-2 infection in human Liver ductal organoids. (**A**) Immunofluorescence staining for SARS-CoV-2 N protein and E-cadherin in human liver ductal organoids. (**B**) Two cases of human liver ductal organoids were harvested at indicated time points following SARS-CoV-2 infection to examine the virus load using qRT-PCR. *RNP* was used as an internal control. Data were presented as mean±s.d. **\*\*\*** indicates *p*<0.001.

### SARS-CoV-2 infection impairs the barrier and bile acid transporting functions of cholangiocytes

The viral load in organoids was significantly decreased at 48 hours post infection (Figure 2B), probably due to virus-induced death of host cholangiocytes or activation of anti-viral response. This promoted us to detect whether SARS-CoV-2 infection could influence the tissue behavior.

The main function of cholangiocytes in homeostasis is to transport bile acid secreted by hepatocytes into bile ducts. The tight junction between cholangiocytes maintains the barrier function of bile ductal epithelium, which is essential for bile acid collection and excretion. We found that SARS-CoV-2 infection ablated the expression of Claudin1 (Figure 3), suggesting that the barrier function of cholangiocytes was disrupted. More importantly, the expression of two major bile acid transporters, apical sodium-dependent bile acid transporter (ASBT) and cystic fibrosis transmembrane conductance regulator (CFTR), was significantly reduced following SARS-CoV-2 infection (Figure 3). These data indicate that SARS-CoV-2 infection impairs the barrier and bile acid transporting functions of cholangiocytes through modulating the expression of genes involved in tight junction formation and bile acid transportation. Our study therefore supports the idea that the liver damage in COVID-19 patients might be resulted from direct cholangiocyte injury and consequent bile acid accumulation induced by viral infection.

**Figure 3.**
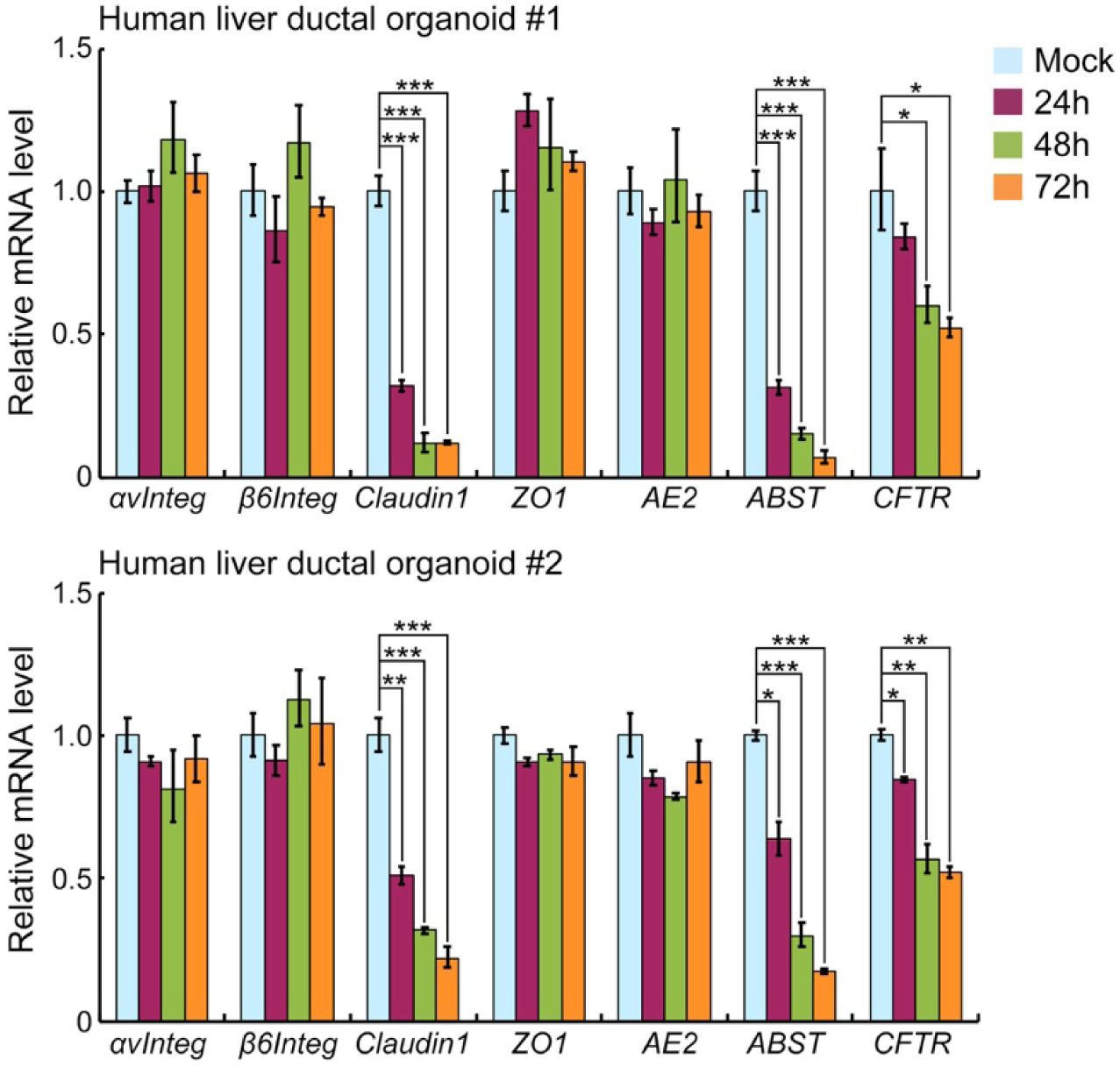
SARS-CoV-2 infection impairs the barrier and bile acid transporting functions of cholangiocytes. Two cases of human liver ductal organoids after SARS-CoV-2 infection were harvested to examine the expression of indicated genes using qRT-PCR. *GAPDH* was used as an internal control. Data were presented as mean±s.d. **\*** indicates *p*<0.05; **\*\*** indicates *p*<0.01; **\*\*\*** indicates *p*<0.001.

## Discussion and Conclusion

Organoids retain the biology of individual tissues, which hold great promise for the study of host–microbe interaction^16^. Here we demonstrated that long-term human liver ductal organoid culture preserves the human specific ACE2+ population of cholangiocytes. The SARS-CoV-2 exposure experiments revealed that the virus infects and replicates efficiently in these organoids. To our knowledge, this is the first SARS-CoV-2-human organoid infection model reported. Given that the culture conditions for various organoids (lung, intestine and kidney) have been established, it would be intriguing to study the tropism, replication, and innate immune response of SARS-CoV-2 infection in other organs that are targeted by this virus.

It appears that liver dysfunction or damage in severe patients with COVID-19 is a common but unignored phenomena. The improper use of anti-viral drugs may be hepatotoxic and cause liver damage. On the other hand, SARS-CoV-2 infection may trigger overwhelming inflammatory response and lead to tissue injury at multi-organ levels including the liver^2^. In this study, by using the human liver ductal organoids as model, we have clearly shown that SARS-CoV-2 can infect the cholangiocytes and impair their barrier and bile acid transporting functions. This could be due to the direct viral cytopathogenic effect on target cells that express the ACE2 as entry receptor. The viral infection may also down-regulate the expression of host genes involved in the formation of tight junction and transportation of bile acid. Thus, it is noteworthy to take into account the fact that the liver damage in COVID-19 patients might be in part the result of direct cholangiocyte injury and consequent bile acid accumulation caused by SARS-CoV-2 infection, which should be cautious in clinical treatment.

By employing human liver ductal organoids, we have investigated the infection and liver tissue damage of SARS-CoV-2 ex vivo. Besides the dissection of viral pathogenesis, this platform could also be applied to evaluate the efficacy of novel anti-viral drugs, especially when ideal animal models are lacking.

## Methods

### Human biopsy

Human liver biopsies were obtained and used for research purposes with approval from the Medical Ethical Council of Zhongshan Hospital. The study abides by the Declaration of Helsinki principles.

### Virus stock preparation

SARS-CoV-2 was isolated from a COVID-19 patient in Shanghai, China (SARS-CoV-2/SH01/human/2020/CHN, GenBank accession no. MT121215). Virus was plaque-purified, propagated in Vero-E6 cells, and stored at –80°C for use. All experiments involving virus infections were done in biosafety level 3 facility strictly following the regulations.

### Liver ductal organoid culture and SARS-CoV-2 infection

The human ductal organoids were generated from primary bile ducts isolated from human liver biopsies as described by Huch et al^17^. The organoids embedded in Matrigel (Corning, 356231) were scrambled off the plate and collected in tubes, then washed with cold PBS by pipetting the material up and down for 10 times. After centrifugation (2 min at 250 g), the organoid pellet was suspended with medium containing 5 μM Y-27632 (Sigma-Aldrich, Y0503). Around 200-300 organoids were infected with 1.2×10^5^ PFU of SARS-CoV-2 in 24 well plate containing 500uL medium and incubated at 37 °C for 1 hour. After incubation, organoids were collected by pipetting and washed once with PBS, then repeated the centrifugation and removed supernatant. The organoids were embedded with Matrigel, followed by seeding on a 24-well plate. After polymerization, culture medium was added.

### Immunofluorescence

For whole mounting liver organoids staining, organoids were fixed in 4% paraformaldehyde for 30 min at 4 °C, washed with PBS and permeabilized with 0.25% Triton X-100 (Sigma-Aldrich, X100) in PBS for 30 min. The organoids were then washed with PBST (PBS containing 0.1% Tween 20) and blocked by 5% BSA in PBST for 1 hour at room temperature. Organoids were incubated with the primary antibodies at 4 °C overnight, washed with PBST 3 times, and incubated with the secondary antibodies and DAPI for 1 hour at room temperature. Organoids imaging was performed on confocal microscope (OLYMPUS, FV3000). The following antibodies were used: rabbit anti-ACE2 (Sino Biological Inc, 10108-RP01, 1:100), rabbit anti-SARS-CoV-2 N protein (Rockland, 200-401-A50, 1:500), mouse anti-E-cadherin (BD Biosciences, 610181), Cy3-conjugated donkey anti-rabbit IgG (Jackson Lab,711-165-152), Alexa Fluor 488-conjugated donkey anti-mouse IgG (Jackson Lab, 715-545-151).

### Quantitative RT-PCR

Total RNA was isolated from organoids by RNeasy Mini kit (QIAGEN,74106) and reverse-transcribed into cDNA with M-MLV Reverse Transcriptase (Invitrogen, 28025013). Quantitative real-time PCR was performed on CFX384 Touch System (Bio Rad). Primers used were listed in Table 1. The SARS-CoV-2 primer and probe sets were provided by Integrated DNA Technologies (IDT, 10006606).

**Table 1.**
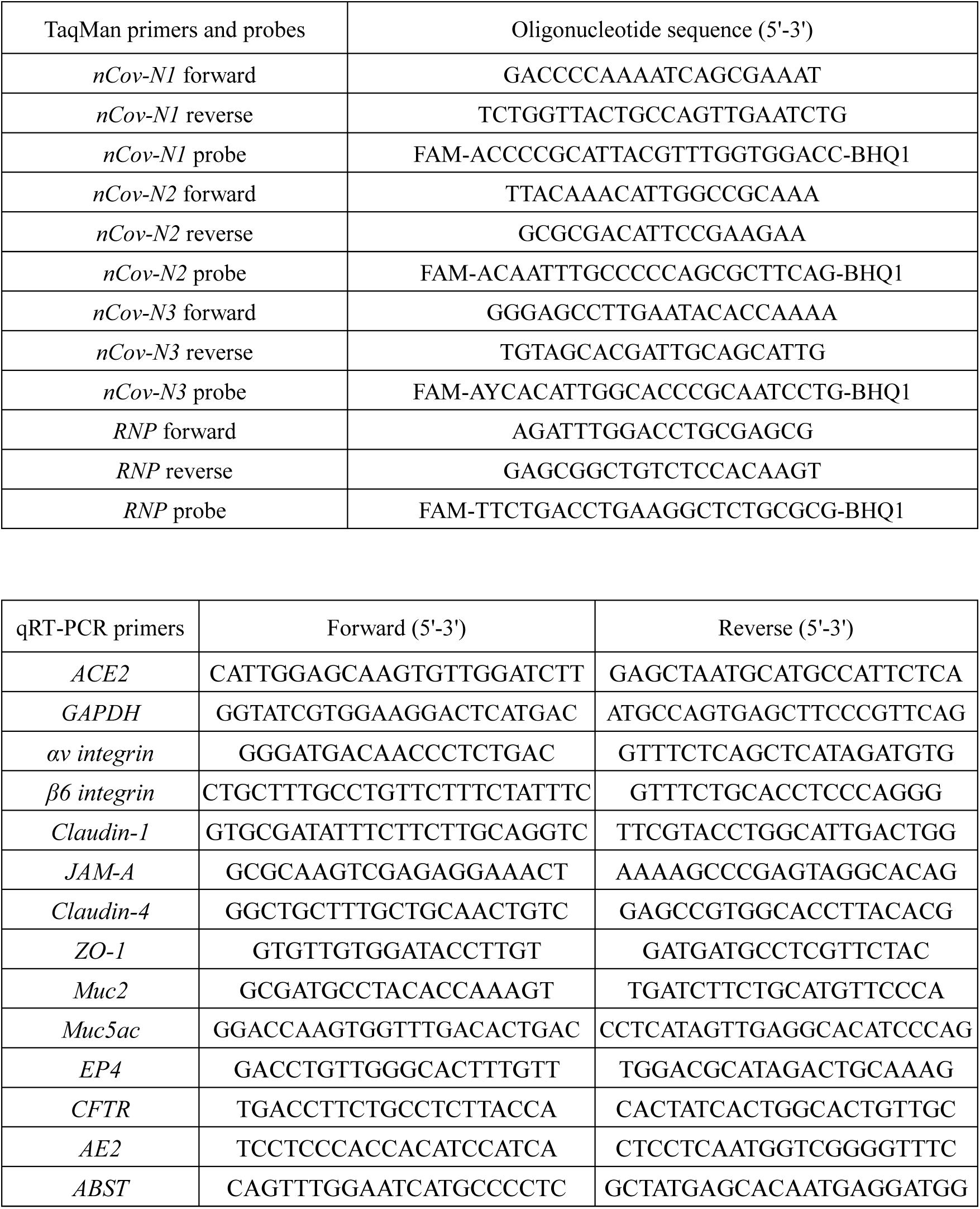
Primers and probes for qRT-PCR.

### Single-cell RNA seq and data analysis

Single-cell RNA sequencing was performed using the 10x Genomics Chromium System. Human liver ductal tissues were derived from a patient who underwent resection, cultured for 3 passages as described above. Mouse primary liver ductal organoids were cultured from biliary ducts isolated from an 8-week-old C57BL/6 mouse. Briefly, organoids were dissociated with 1× TrypLE Select Enzyme (Gibco, 12563011) to obtain single cell suspension. A total of around 8,000 cells per sample were captured on a 10×Chromium device and library preparation was carried out using Single Cell 3’ Reagent Kits v2 according to the manufacturer’s instructions (10× Genomics). Libraries were sequenced on an Illumina NovaSeq 6000 platform.

Cell Ranger (version 3.1) with default parameters was used to process sequencing data to generate feature-barcode matrices. The human dataset was analyzed using the standard workflow on the Seurat R Package (version 3.1.3) (Butler et al., 2018). For the feature-barcode matrix of 8,094 cells from the human dataset, we removed cells with less than 500 genes and more than 6,000 genes as well as cells with high fraction of mitochondrial UMIs (> 20%). 7,978 high quality cells and 17,447 expressed genes were remained for downstream analysis. The cell populations were clustered using the ‘FindClusters’ function and visualized in 2 dimensions by t-distributed stochastic neighbor embedding (t-SNE) derived from the top 10 principal components. For the feature-barcode matrix of 9,690 cells from the mouse dataset, we retained cells with expressed genes between 500 and 6,000 as well as cells with low fraction of mitochondrial UMIs (< 10%). Finally, 8,812 high quality cells and 16,019 expressed genes were remained for downstream analysis. The integration of human and mouse datasets was processed by the standard Seurat v3 integration workflow.

### Statistical analysis

We employed Student’s *t*-test or ANOVA test to analyze the parametric experimental results. Significant differences were noted with asterisks.

## Acknowledgments

The authors thank Dr. Stacey S. Huppert for technical assistance. We also wish to acknowledge Di Qu, Xia Cai, Zhiping Sun, Wendong Han and the others at Biosafety Level 3 Laboratory of Fudan University for experiment design and expert technical assistance. This work was supported by grants from the National Key Research and Development Program of China (2018YFA0109400 and 2018YFA0109800), the Zhejiang University Special Scientific Research Fund for COVID-19 Prevention and Control (2020XGZX013) and the Shanghai Municipal Science and Technology Major Project (2017SHZDZX01). B.Z. was sponsored by Shanghai Rising-Star Program and Eastern Scholar Program.

## Author Contributions

B.Z., C.N. and R.Z. conceived the study; B.Z., C.N., R.G., Y.W., L.Y., J.W., T.L., J.L., W.X.,. and R.Z. performed the experiments; B.Z., J.L., R.Z. and X.L. supervised the work; Y.X X.W. and Z.Y. contributed to the discussion of the results; and B.Z., C.N., R.Z. and X.L. wrote the manuscript.

